# American Crow responses to habitat desaturation by West Nile virus

**DOI:** 10.1101/2024.02.01.578463

**Authors:** Carolee Caffrey

**Author notes:** Cite as: Caffrey, C. 2017. American Crow Responses to Habitat Desaturation by West Nile Virus. ACB. www.caroleecaffrey.com.

## Abstract

In late Summer 2002, West Nile virus spread to a population of individually-marked, cooperatively-breeding American Crows (*Corvus brachyrhynchos brachyrhynchos*) in Stillwater, OK. Within six weeks, approximately 42% of adults were dead, leaving widows, widowers, and vacant territories. I looked to see if surviving unpaired adult crows left groups to occupy vacant territories, as predicted by theory (Habitat Saturation/Ecological Constraints Hypothesis; Emlen 1982, 1984). Survivors did not behave as predicted, had previous decisions to delay breeding and live in others’ groups been made in response to a saturated habitat. Their aberrant behavior and physical attitudes suggested their losses and grief affected them in ways not included in simplistic models of avian behavior.

## Introduction

In mid August 2002, the resident population of American Crows (*Corvus brachyrhynchos brachyrhynchos*) in Stillwater, OK, had been individually marked and under observation for five years. Groups of up to ten individuals, including pairs and auxiliaries of both sexes, various ages, and diverse dispersal histories (Table 1), occupied all-purpose territories distributed throughout town (Fig. 1). By later in the month, West Nile virus (WNv) was detected in Oklahoma - approximately 130 km east of Stillwater - and by mid September a marked crow was dead. By the end of October, approximately 42% of adult population members had fallen victim to the virus (Table 1; Caffrey et al. 2003, 2005), leaving widows, widowers, and vacant territories. The situation appeared to offer a naturally-occurring, direct test of the Habitat Saturation context of the Ecological Constraints Hypothesis (ECH), formulated by Emlen (1982, 1984), to explain decisions by auxiliaries in cooperatively-breeding species to delay breeding and dispersal.

**Table 1.**
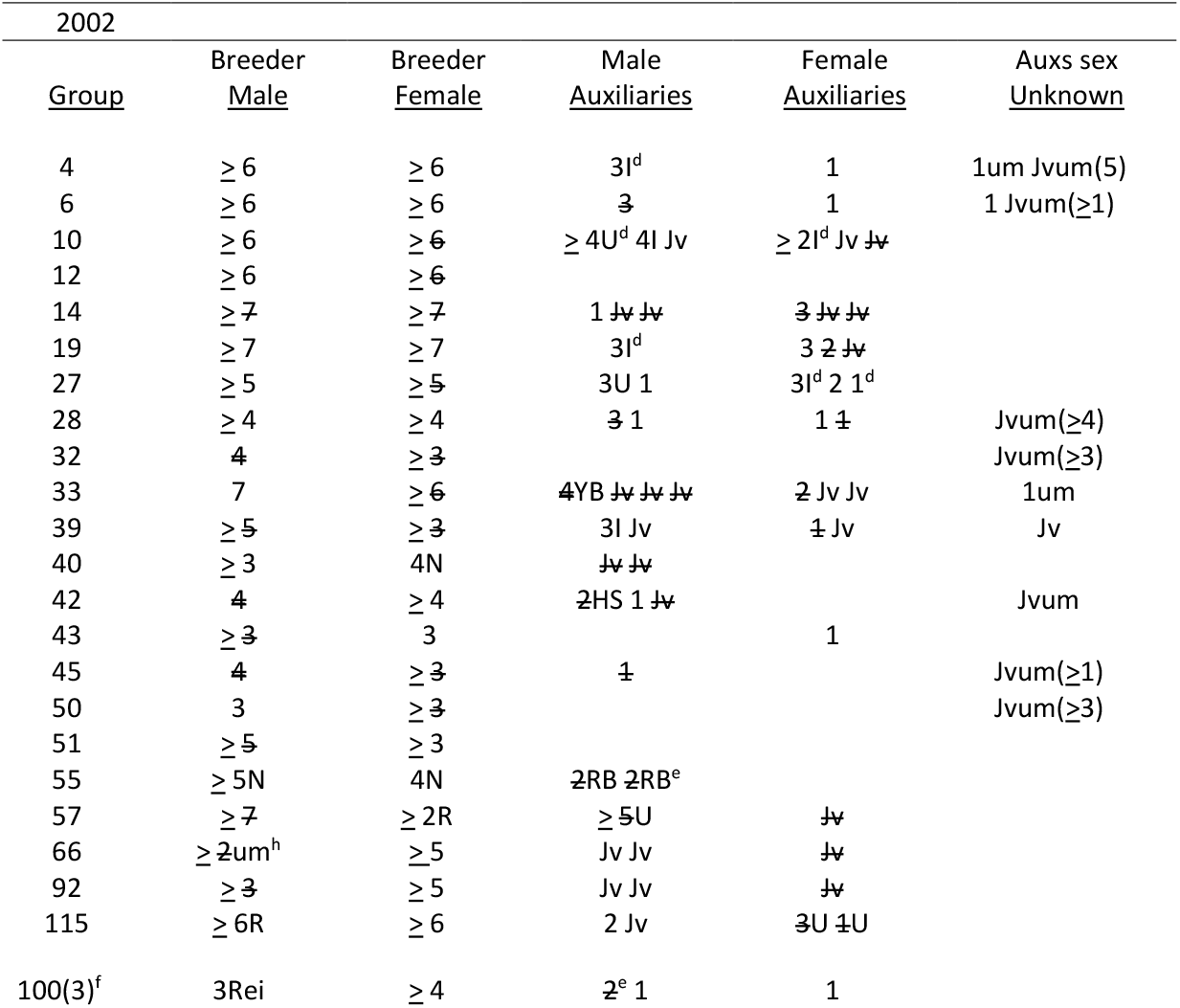
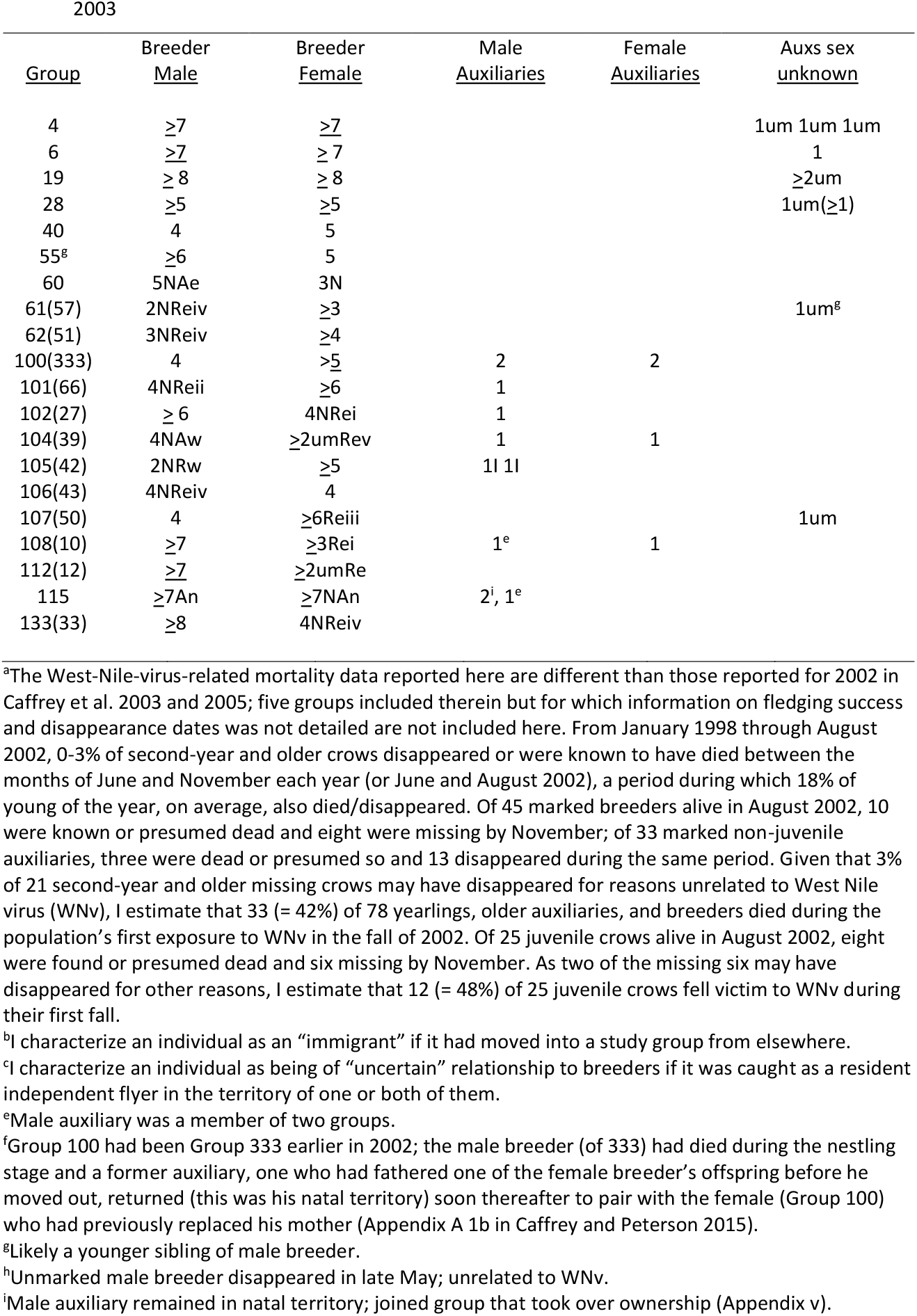
Composition and fate of members of pre-hatch groups of American Crows in Stillwater OK, USA (Caffrey et al. 2015) and young of the year surviving to fall in 2002, and composition of pre-hatch groups in 2003. Each numeral represents an individual by its age in years; Jv (juvenile) indicates young of the year. Strikeouts indicate individuals found dead, presumed dead, or who disappeared during September and October 2002^a^. N (novice): known to be breeding for first time. A: assumed ownership of vacated territory; w: auxiliary from within group (originally immigrant), e: extragroup auxiliary budded/moved in as original pair shifted (Appendix iv), n: neighboring pair shifted to take over part of territory (Appendix iv and v). R: breeder was replacement for territory owner’s mate during preceding nesting season; in 2003, w: from within group (offspring from previous year still in natal territory; Appendix iii), e: extragroup individual (i: previous group member returns [LH, Appendix A 1b in Caffrey and Peterson 2015, and RM, Appendix A 2d in Caffrey and Peterson 2015], ii: offspring from previous year returns [Appendix vi], iii: widowed breeder from different group [Appendix i], iv: auxiliary from different group, v: unmarked individual of unknown history replaces breeder of same sex to pair with new territory owner). Most auxiliaries had hatched in nests of one or both breeders. I: immigrant^b^. HS: social half-sib of male breeder (Appendix iii; EK, Appendix A 2j in Caffrey and Peterson 2015). YB: younger brother of male breeder (KR, Appendix A 2a in Caffrey and Peterson 2015). RB: raised by male breeder when in previous group; one is son (an occasion of reproductive sharing), one not closely related (Appendix A 1b, and Appendix B Predictions 7 and 8 [#1], and 12-15, in Caffrey and Peterson 2015). ^d^: dispersed at hatching (Caffrey et al. 2015); not member of group in Fall. U: dispersal history (relative to group membership) uncertain^c^. um: unmarked (#s of unmarked juveniles in parentheses).

**Figure 1.**
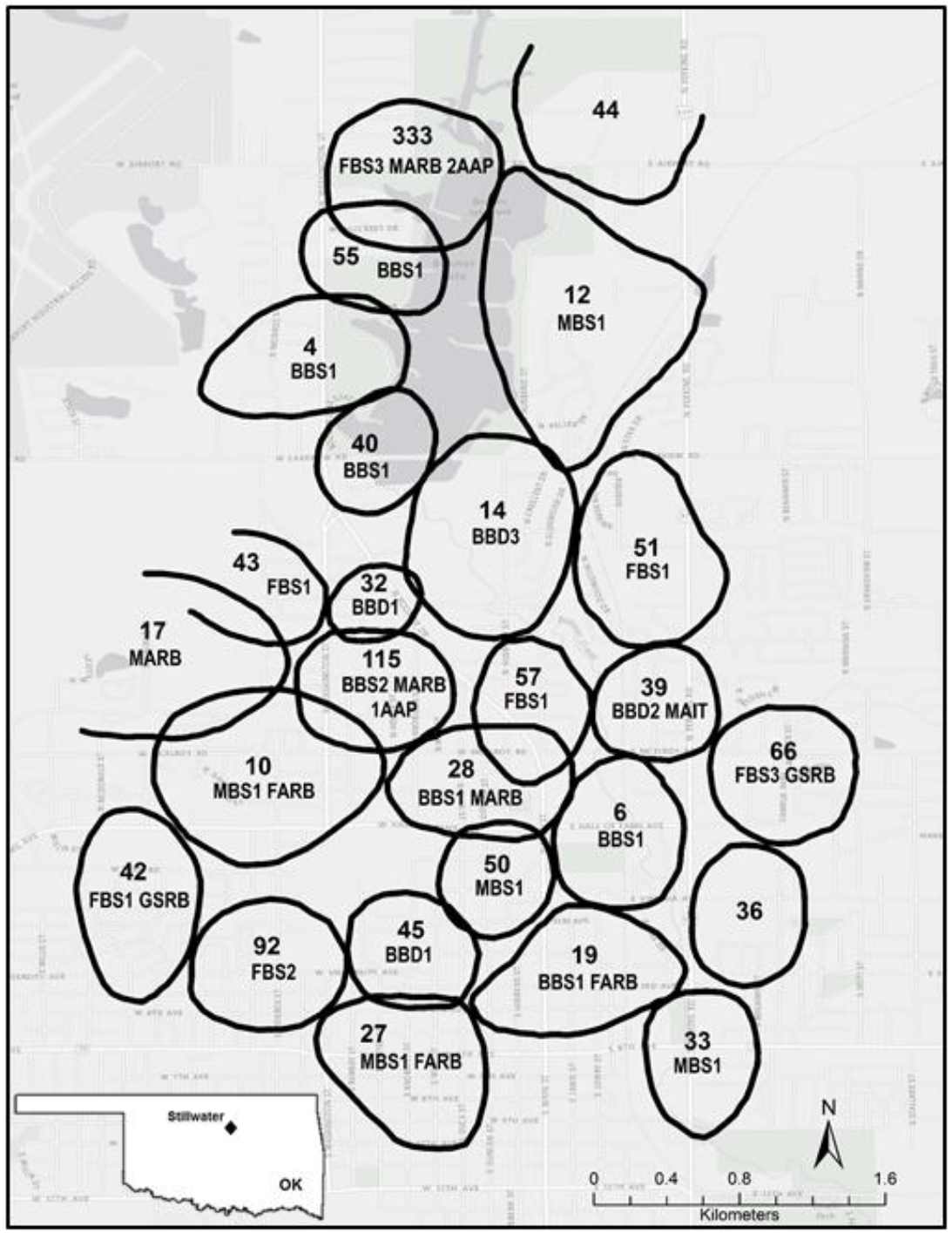
Approximation of the territories of study groups of crows in Stillwater, OK in August 2002. BBS (both breeders survive): 1 - remain in territory, 2 - shift to another territory. BBD (both breeders die): 1 - territory vacant in 2003, 2 - auxiliary inherited, 3 - different pair moves into parts. FBS (female breeder survives): 1 - remains in territory/male replaced, 2 - moves to different territory/territory remains vacant in 2003, 3 - male breeder disappeared earlier in year; replaced as breeder by previous auxiliary (Group 333) or by son of female (66). MBS (male breeder survives): 1 - remains in territory/female replaced. FARB: female auxiliary replaced dead breeder (one in a different group, and two returned to former groups). MARB: male auxiliary replaced dead breeder (three in different groups; one had replaced male earlier in year). 1AAP: one adult auxiliary present in 2003; 2AAP: two adult auxiliaries present in 2003. GSRB: genetic son of female replaced male breeder. MAIT: male auxiliary inherited territory.

The ECH has itself evolved over the years so as to also incorporate possible benefits available through philopatry (as envisioned by Stacey and Ligon 1987, 1991, Wasser 1988, Zack 1990), yet at its core, since its inception and early iterations (Selander 1964 [in Brown 1978 and Koenig et al. 1992], Brown 1969, and e.g., Verbeek 1973, Gaston 1978, Stacey 1979), has been the idea that for many species, nonbreeders associating with breeding pairs during nestling seasons have chosen to do so because of a lack of “suitable” breeding space (given their first choice would be to breed on their own). Over years subsequent to the theory’s formulation, positive associations between scarcity of suitable habitat and delayed breeding have been demonstrated for many species through long-term observational studies of natural patterns and experimental manipulation of breeding opportunities, including the removal of breeders (e.g., Walters 1991, Hatchwell 2009 and references therein, Hatchwell and Komdeur 2000 and references therein).

I looked to see if surviving unpaired adult crows left groups to occupy vacant territories, as predicted by theory, if lack of suitable habitat had been an important part of previous decisions to not breed independently. The aberrant behavior and physical attitudes of survivors suggested that their loss of family and friends, and the shattering of their social lives and collective crow umwelt, affected them in ways not included in simplistic models of avian behavior.

## Methods

This crow population had been under observation since Fall 1997. Individual crows had either been obtained as nestlings or caught as free-flyers (Caffrey 2002a), and were marked with patagial tags and colored leg bands (Caffrey 2002b, 2002c). From blood samples, the sexes of all individuals were determined using the 2550/2718 primer set (Fridolfsson and Ellegren 1999), parentage was analyzed in the program Cervus (Kalinowski et al. 2007), and microsatellite-based relatedness coefficients (r) estimated in the program Relatedness (Queller and Goodnight 1989); details in Caffrey and Peterson (2015). Crows are thought to not breed until their third calendar year at the earliest (Black 1941, Townsend et al. 2009), and they molt to adult plumage half-way through their second year (Emlen 1936); until then, immature individuals could easily be distinguished from adults. Thereafter, free-flyers were initially assigned the age ≥2 years when caught.

During 2002-2003, Stillwater was a town of approximately 40,000 people within a 72.51 km^2^ area (U.S. Census Bureau). Crows based their activities throughout the year in all-purpose territories (50-150 ha in size) distributed throughout public, commercial, and residential areas (Fig. 1). Observations of crows were made with binoculars and spotting scopes, primarily from vehicles.

Many auxiliary crows in Stillwater dispersed out of groups each year as hatching was imminent (Caffrey and Peterson 2015), and so nesting-season group composition for this population has been described in pre- and post-hatch contexts (Caffrey and Peterson 2015, Caffrey et al. 2016a, 2016b). Had lack of appropriate space been influencing crow decisions to delay breeding, relaxation of the constraint would affect establishment of territory ownership, and so I follow survivor responses to space being made available in 2002 only through pre-hatch group composition in 2003 (Table 1).

Once WNv reached Stillwater, many crows fell victim (Caffrey et al. 2003, 2005). A crow was classified as dead if its carcass was found or it was seen staggering, unable to balance, or inactive and appearing sick prior to its disappearance. One was classified as presumed dead if its disappearance co-occurred with having juveniles present and/or palpable differences in post-disappearance behavior of remaining group members (Discussion).

## Background

In 2002, the pre-hatch population of marked crows comprised 29 groups, 26 for which my students and I knew the histories of many individuals (Fig. 1), and 23 for which we had robust group composition data (Table 1; two unsuccessful first attempts wherein group size was 6 [10a and 115a; Table 1 in Caffrey and Peterson 2015] not included here). Group size varied from 2 to 5, and many groups contained female and male auxiliaries of different ages, dispersal histories, and relationships with breeders (Table 1), including – across the population - 8 and 15, respectively, third-year and older females and males (Table 1). In mid August, subsequent to dispersal of six auxiliaries (Table 1) and the early post-fledging deaths of some young of the year, group size ranged from 2 to 10, including – across the population - 14 and 18 marked female and male, respectively, auxiliaries old enough to breed in (upcoming) 2003 (Table 1). Within a month, WNv was detected in Oklahoma and a marked crow was dead (Caffrey et al. 2003). By the end of October, eight more carcasses of population members had been found (Caffrey et al. 2003), and an estimated 42% of marked second-year and older crows had fallen victim (Table 1). Victims included 18 breeders, leaving four widows and six widowers (Table 1), and four territories without current owners (Fig. 1).

## Results

The first two crows to replace lost breeders were females who had previously lived in the groups of surviving widowers (Groups 102 and 108, Table 1); both females, originally immigrants, had moved out earlier in 2002 (Groups 27 and 10, Table 1), and both returned within 1-4 days of female breeder deaths. Prospectors and ultimate replacements continued to be seen with surviving breeders soon after mate losses such that by late November, all widows and widowers had paired with whom they would mate in 2003 (Table 1). All but two other replacement mates of known history had been auxiliaries in other study groups; the exceptions were a widow and widower who paired with each other: the female moved out of and vacated her territory (#92, Fig. 1) to move in with the male (#50, Table 1 and Fig. 1; creating Group 107 [Table 1, and Appendix i]; her two surviving juvenile sons stayed together through the process of incorporating themselves into a next-territory group, then ended up manifesting different types of relationships with the pair; Appendix ii).

Four territories were rendered owner-less upon the deaths of both breeders (#s 14, 32, 39, and 45; Fig. 1), and one territory was vacated upon the widow’s move to pair with a widower in his territory (92 and 50, respectively, Fig. 1, above, and Appendix i). Territory 39 was “inherited” by a resident adult male auxiliary (originally an immigrant), who paired with an unmarked female of unknown history. Parts of Territory 14 were taken over as Group 115 shifted north and east (leaving their previous space to a new pair; below). Large areas associated with successful crow nesting in the past, including Territories 32, 45, and 92, and parts of 14, remained unoccupied in 2003.

In addition to the above, one new group (#60; Table 1) formed early in the nesting season of 2003 as two novices paired and settled in the space previously occupied by Group 115 (above, and Appendix iv). Four adult crows, three of them marked (one female and two males), chose to live as auxiliaries in study groups in 2003 (Table 1 and Fig. 1).

## Discussion

Crows did not behave as predicted had previous decisions to delay breeding and live in others’ groups been largely a function of a saturated habitat, but the situation turned out to be more than just a matter of space becoming available to would-be breeders kept in check by the lack thereof. WNv destroyed the underpinnings of the fabric of crow society, a society based on high survivorship and long-term social relationships wherein cooperation and planning for the future occurred. Neighborhoods and communities had included several generations of several families, the individuals of which lived together, with their own families, or with friends. Neighbors and community members had foraged, loafed, and warned of and harassed predators together. WNv’s arrival created a situation these organisms had never encountered before, and upended selection pressures presumably in place for a long time. It is true that space capable of supporting successful breeding was now widely available, but the neighborhoods and community of support and protection were gone. What becomes most important then? As far as I know, no one has articulated a complex-enough social theory to make predictions about how individual animals should respond to catastrophic population loss.

A clear pattern emerged, though, as breeders began to die: opportunities for previously unpaired crows to pair with (known) surviving breeders in known territories were seized upon almost immediately. Of four male replacement breeders, two had lived next territory (Groups 61 and 106, Table 1), one had lived two territories away (Group 62, Table 1), and one was the genetic son of the surviving widow (Appendix iii). Of six replacement female breeders, one had lived next territory (Group 133, Table 1), two had been former auxiliaries (Groups 102 and 108, Table 1), one had been a breeder two territories away (Group 107, Table 1), and two were unmarked. The two unmarked replacements occurred at the eastern edge of my study population (Groups 104 and 112; 39 and 12 in Fig. 1), where groups were close neighbors with many unmarked individuals. In addition, the auxiliary who acceded to male breeder in Group 104 (formerly Group 39, Table 1), had been seen several times near the beginning of the 2002 nesting season with an unmarked individual – once pulling on twigs (a part of the nest-building process) – before the unmarked crow was no longer present as Group 39 began nesting. Thus it is probable that both of the unmarked replacements were already familiar with their new mates and territories, too.

The single pair that formed anew between crows not already living in the space that was to become their territory included a male who had lived next territory for several years and had also recently spent a lot of time there. The space itself was made available through a (surviving) pair’s shift, and not through death or departure via WNv (Appendix iv). (The pair that shifted *did* move into space opened up in such a way, though - parts of Territory 14; Fig. 1; the only such space to be reoccupied.)

Other parts of Territory 14, and the three other territories available through WNv-related death/departure - 32, 45, and 92 (Fig. 1) - remained unoccupied in 2003. Four adults chose to live as auxiliaries rather than attempt to breed on their own that year (Table 1), but the three of them that were marked were only two years old (only one crow in five years had attempted to breed at that age; below). In addition, these crows had witnessed the inexplicable deaths of all or most of the members of groups in vacated territories; individuals that included relatives and friends (Caffrey and Peterson 2015). Crows are known to avoid areas associated with conspecific misery and distress (hence the somewhat successful use of crow carcasses and effigies to dissuade live crows from entering seabird colonies or roosting in particular areas; e.g., Peterson and Colwell 2014 and Avery et al. 2008, and personal experience). In another population similarly affected by the arrival of WNv, crows in Ithaca NY only very slowly moved into space vacated by the deaths of most group members (over years; Clark et al. 2006). The misery and distress surviving crows in Stillwater had observed and experienced was intense, and so it is possible that areas vacated in Fall 2002 were actively avoided in Spring 2003 (the “haunted house” effect; A Clark, pers. comm.). AM’s abandoning her territory after the death of her mate, a juvenile, and two suitors (Appendix i) supports such an idea; Group 115’s shift into parts of Territory 14 does not.

Decisions to delay breeding in 2003 may also have been related to mitigation of grief (below); one of three marked adult auxiliaries in 2003 had been the sole survivor of a large family (he joined the next-territory group that shifted to claim part of his parents’ territory; Appendix v). The two others were siblings who spent much of their time through the end of 2002 and into the nesting season of 2003 in close proximity to their mother and social-”uncle”-now-stepfather (Appendix vii). Choosing to delay breeding and continue to live with and among co-surviving relatives and friends in the short term may have been adaptive over the long term - young feral horses that lost both parents and all other group members had higher survivorship when in networks with greater numbers of associates (Nuñez et al. 2015) – except that all three marked adult auxiliaries, and all others but one in their groups, fell victim to WNv in 2003 (the second, more devastating, year of WNv presence in Stillwater; Caffrey et al. 2005). Young Carrion Crows (*C. corone*) exhibit greater philopatry in resource (food)-enhanced territories (Baglione et al. 2006); could it be that social and emotional support are also resources upon which crows depend?

That higher vertebrates, including humans, share among them many aspects of structural and physiological characteristics, including those of brains, cannot be disputed. That at least some of those other higher vertebrates share with us the capacity to feel emotions has been postulated in the scientific literature since at least the time of Darwin (1859, 1871, 1872); recent evidence leaves no doubt (e.g., Griffin 1981, Bekoff 2000a and references therein, Bekoff 2000b and references therein, Ristau 2016). Members of the genus *Corvus* share with humans some extraordinary cognitive capabilities (e.g., Emery, 2006 and references therein, Loretto et al. 2012 and references therein), and American Crows live complex social lives involving relationships with others that endure for years (Caffrey and Peterson 2015). Although nonhuman feelings and emotions are not going to be exactly the same as human ones, using language to describe them – words such as “want,” “pleasure,” and “grief” – enables such experiences, through an animal’s behavior, to be accessible to us (Griffin 1981, Bekoff 2000a). People often agree on what they think an animal is likely feeling (Bekoff 2000a), and Heinrich (1999) used the word “love” in the context of ravens paired for years (without quotation marks; p. 341). Grief is identified by comparing the behavior of survivors before and after deaths of mates, relatives, or friends; differences in social behavior and expression of emotional affect may persist for weeks (King 2013; through personal observation, I suspect much longer). Grief has been identified in elephants (*Loxodonta africana;* e.g., Poole 2000), chimpanzees (*Pan troglodytes*; e.g., Goodall 2000), orcas (*Orcinus orca;* Rose 2000), dolphins (*Stenella frontalis*; Herzing 2000), baboons (*Papio Anubis;* e.g., Smuts 2000), and coyotes (*Canis latrans;* Bekoff 2000a); all social species wherein individuals are tightly bound to each others’ lives. My students and I saw sad and grieving crows, and our hearts broke for, and with, them.

As crows began to die, my students and I saw differences in the postures and movements of surviving group members; at times they seemed to walk around territories aimlessly… Were they truly moving more slowly? Were their heads and shoulders actually drooped? It is difficult to pinpoint the aspect of them that had changed, but it was palpable enough to warrant, in field notes, an occasional inclusion of something akin to “All seems almost normal today.” Crows never before seen together were seen together. Orphaned juveniles and yearlings (in 2003) produced gut-wrenchingly sad and plaintive vocalizations, bringing to mind Skutch’s (1996) observation that bird sounds seem to parallel their feelings in ways similar to our own.

In addition, in the weeks of deaths between mid-September and late October, and then continuing into November, interactions between neighboring groups, especially those wherein breeders were being replaced, were more overtly aggressive than had ever been observed previously (see Appendix B in Caffrey and Peterson 2015 for details of relevant behavior). Crows chased and vocalized at each sometimes several times a day, as territory boundaries were maintained or shifted slightly, or one flew through another’s area (not a cause for alarm in the past; Caffrey and Peterson 2015); they seemed “on edge.” Three groups contested space opened up by the death of the breeders and all but one auxiliary of Group 14 (Fig. 1, Appendix v), and the contests seemed particularly highly charged. Juveniles of surviving breeders acted submissively toward possible and ultimate replacements of both sexes, some of whom behaved very aggressively toward them. Possibly the most aggressive of all – toward the three juveniles present also - was a second-year male (RZ) competing with a third-year male (Appendix i) to replace a breeder whose widow was smart (AM, in Appendix 4 in Caffrey et al. 2016b), savvy (Appendix 3g in Caffrey et al. 2016b), an extraordinary parent (Fig. 1 in Caffrey et al. 2016b), and reproductively successful. In humans, heightened aggression is a common reaction to many forms of trauma (e.g., Kivisto et al. 2009 and references therein, and Bell and Orcutt 2009 and references therein).

Grief and responses to trauma may also have had something to do with the pairing of two widows with their sons; something never known to have happened before. Townsend et al. (2010) mention occasional mother-son incest in a population of crows in New York, and offer possible adaptive explanations, yet such incestuous extra-pair matings do not pertain, as mothers and sons in Stillwater behaved as social pairs in addition to being the genetic parents of the young in their nests. One of the two males had never left his natal territory (Group 42 -> Group 105, Table 1 and Appendix iii); the other (Group 66 -> 101, Table 1), after having moved out for 1.5 years, returned soon after his mother’s mate disappeared (earlier in 2002 [May] and not related to WNv; Appendix vi). Neither female had been observed being visited by potential suitors, through winter and early spring, and both pairs looked “normal” during successful nesting attempts in 2003. Such pairings have never occurred among study crows in Ithaca NY, either (K. McGowan, pers. comm.); presumably short-term emotional responses and/or grief mediation overrode the long-term avoidance mechanisms of four of them in Stillwater in 2002-2003.

Possible haunted houses and grief notwithstanding, the ways in which unpaired crows had established themselves as breeders in past years also suggested that lack of suitable habitat was not the basis of decisions to delay breeding. Each year, in addition to the presence of many unpaired adults in my study population (Caffrey and Peterson 2015) was unused space that appeared suitable for nesting and living, and in fact, over subsequent years, would eventually become occupied. Budding occurred regularly – at least eight times prior to August 2002; five for which details of the crows involved and outcomes of nesting attempts were known. All cases involved no obvious aggression from neighbors; in several of them, neighbors appeared to shift territory boundaries to accommodate budders. Auxiliaries who budded from their groups included two genetic sons of both breeders (DY in Group 32 budded from Group 14, and Group 50 [OV] from Group 6; Table 1 and Fig. 1), a male immigrant (HE in Group 45, from Group 19; Table 1, Fig. 1, and Appendix A 2e in Caffrey and Peterson 2015), a male of unknown dispersal history regarding the group from which he budded (NX in Appendix A 1b in Caffrey and Peterson 2015; Group 333 to Group 55; Table 1 and Fig. 1).), and a female of the same status (TF: Group 12 to 40, Table 1, Fig. 1, and Appendix A 2i in Caffrey and Peterson 2015).

The budding of territories by females in a population of crows in Ithaca, NY, had been “very rare” (Clark et al. 2006), so much so that the spike in its occurrence following a Fall of high WNv-related mortality was included under “unusual social changes” (Clark et al. 2006). The female budder in Stillwater was accompanied by a male who had immigrated into her group (Appendix A 2i in Caffrey and Peterson 2015); as two of seven other budders in Stillwater were immigrant males, it is possible that TF’s budding was driven by her mate. He had remained with Pair 12 the previous nesting season (whereas TF appeared to be forced out; Appendix A 2i in Caffrey and Peterson 2015), and so possibly had “earned the right” to bud - if such a mechanism existed. Yet the pair budded into space owned by Pair 4 and not Pair 12 (Fig. 1); a pair with whom TF had long interacted (her mate had been unmarked and so any possible relationship with Pair 4 was unknown). Given the equanimity exhibited by members of this population (e.g., Caffrey and Peterson 2015), I would not presume TF was not integral to the budding process.

One member of each budding pair was known to be breeding for the first time (including two females [TF, and NX’s mate]). Based on mouth color patterns at marking (Emlen 1936), it is very likely that at least three others were also novice breeders (and so at least eight of 10 individuals). Seven of them were known to have previously assisted in others’ nesting attempts. Four of five budding events resulted in fledged young in the first year; the single failure occurred late in the nestling stage and was attributable to a heavy storm (the male of this pair [OV; Group 50 in 2002, Table 1] was the only second-year crow in Stillwater to attempt to breed independently). Two of the successes were those of second attempts (DY, NX) but two were those of first (TF, HE); HE’s “success,” however, was a single fledged male fathered by someone other than him (Appendix A 2e in Caffrey and Peterson 2015).

And so it was possible for two novice breeders to nest successfully together. In addition, successful breeding in Stillwater did not require much space; Group 115 (Fig.1, and Appendix iv and v), e.g., regularly fledged young who survived to adulthood. As such, and all else being equal, adult crows should have been attempting to breed on their own, rather than spending years in others’ groups (Emlen 1982, 1984). What might it have been, then, that made it more profitable for them to delay breeding before WNv struck, and, in fact, thereafter as well?

Acknowledged since the early 1990s is the understanding that maturing individuals in cooperatively-breeding species will assess the consequences of different behavioral options and subsequently behave in ways that maximize lifetime fitness (e.g., Emlen 1991, 1994, Walters 1991, Koening et al. 1992, Komdeur 1992); if opportunities of worth are available, individuals should disperse and attempt to breed (Plans A); if not, Plans B should be about maximizing benefits over lifetimes. It is difficult to imagine the cost/benefit conditions that favored delaying breeding and possibly contributing to the reproductive success of unrelated individuals (Caffrey et al. 2016b) over pairing with a friend and attempting to raise offspring in space available in the neighborhood, but Brown Jays (*Cyanocorax morio*), too, delay breeding and remain in groups when breeding habitat is available (Williams et al. 1994; the authors attribute the behavior to intrinsic benefits of sociality). Crows in Stillwater benefitted in many possible ways from group living (Caffrey and Peterson 2015), but their delaying breeding appeared to also have had something to do with not “being ready,” or – apparently - not being able to pair with someone else who was: adult auxiliaries of both sexes almost immediately dispersed to pair with known, experienced breeders in territories with which both mates would be familiar. That dispersing auxiliaries were not dissuaded by openings becoming available through the presumably scary and confusing population-wide agent of death suggests such opportunities were of great worth.

Environmental conditions (both weather- and human-related) were variable in Stillwater, and many individual crows moved into and out of two or more groups in the years before breeding (Caffrey and Peterson 2015). Caffrey and Peterson (2015) speculated that crows delayed breeding, despite available space and potential mates, as they chose to “add skills, networks of social relationships, and savvy to their toolboxes.” Unfortunately for this population of crows, the factors influencing the short-term decisions of individuals pursuing long-term plans will forever remain unknown.

## Acknowledgments

I wish to first thank Shauna Robertson Smith and Tiffany Weston Hackler, my graduate students at the time, for continuing to press on in the field under extremely emotionally-draining circumstances. I wish to next thank those who helped me manage my own emotions and think clearly about what could be learned from the situation, especially Kevin McGowan, Charlie Peterson, and Anne Clark.

As this study was possible only after years of catching, marking, and watching the crows of Stillwater, I thank, again, all of the people and organizations acknowledged in Caffrey and Peterson 2015, and Caffrey et al. 2016 a and b. For this manuscript, I am extremely grateful to Morgan Wehtje for Figure 1, Marc Bekoff and Carolyn Ristau for information regarding animal grief, and Charlie Peterson and Fritz Hertel for helpful editorial comments on a draft.

This work was supported by an NSF SGER award (0331531).

## Appendix

i. AM, the female breeder of Group 92 in 2002, lost her mate to West Nile virus (after reuniting with him the previous year; Appendix A 1a in Caffrey and Peterson 2015) by late September. Within days, two male auxiliaries from neighboring territories began visiting with AM and acting aggressively toward each other; one (RZ; the son of a female breeder and male auxiliary from different groups [Appendix A 2e in Caffrey and Peterson 2015] notably so; he aggressively attacked and chased AM’s three juveniles also. Both suitors and one of AM’s juveniles disappeared by mid October. A few times in the subsequent two weeks, AM was seen in places other than her territory. Days after the female breeder of another group (# 50; Fig. 1) was seen staggering and then disappeared, AM was observed with the widowed breeder (OV) in an area near his territory. Through November, she was seen increasingly frequently with him before moving in.
ii. AM’s two surviving, very similarly-sized juvenile sons remained attached to their natal territory (#92, Fig 1) as she began to prospect in mid-late October and were seen with the female breeder of Group 45 (to their east; Fig. 1) before her death. They remained pretty much alone together in natal areas through mid November, when they were seen with a widow (YL) and her one-year old son (Group 42; below, iii) in an area between their two territories (Fig. 1); one of AM’s juveniles (FC), but not the other, was acting submissively to, and being aggressed upon by, YL. The two of them moved in with YL and her son over the next month, although FC did so completely and the other only part-time, for about a month, until he was no longer seen in Stillwater. Clearly the values for the variables involved in cost/benefit analyses regarding what to do next differed for these two same-sexed broodmates.
iii. Group 42 in August 2002 consisted of a pair, the male breeder’s two-year old half-brother, and the pair’s one-year old son (SN) and two juveniles (Table 1, and Appendix A 1a and 2j in Caffrey and Peterson 2015). The male breeder was one of our first population members to die from exposure to WNv, in mid September; his and his mate’s two juveniles died soon thereafter. His half-brother seemed in the process of pairing with his widow (YL) until he died, too, in mid October (Appendix A 2j in Caffrey and Peterson 2015). In mid November, two juvenile males from a neighboring territory began to move in with YL and SN (above, ii). One of them (FC) remained through the following summer. An unmarked yearling joined the group in January through March 2003, with no observed aggression involved (YL was observed preening both FC and the unmarked individual in March). The unmarked individual was no longer around as YL and SN began behaving as a pair and working on a nest. FC was observed contributing to both nest building (rare in auxiliaries; Caffrey et al. 2016a) and nestling feeding. Mother and son fledged three young. YL and one of the three were dead by the end of June, SN and another by mid September, the third by late September, and FC by mid October.
iv. TM had been an auxiliary in Group 10 since early 1999 (he moved in from next-territory when a yearling; Appendix A 2d in Caffrey and Peterson 2015), and had been interacting regularly with members of Group 115 (Fig. 1) since then. Upon Territory 14 becoming ownerless in the Fall of 2002, Group 115 began competing for and utilizing parts of it, while continuing to remain attached to their already-owned space. As the nesting season of 2003 got underway, TM was repeatedly observed in Territory 115 (at times co-occurring with group members, with no obvious signs of distress) and in what had been the western end of Territory 28 (Fig. 1; Group 28 shifted slightly east as Group 39 [now 104] shifted slightly north). By mid March, Group 115 had shifted and settled into the southwestern section of what had been Territory 14, and TM paired and nested in what had been Territory 115. TM’s mate - DX - had previously moved out of and back into her natal territory (#27, Fig. 1), but had recently been observed in several different locations around town, following the WNv-related death of her mother the preceding Fall (Appendix A 2d in Caffrey and Peterson 2015). DX was *not* one of the one or two females with whom TM had built two previous nests (Appendix A 2d in Caffrey and Peterson 2015).
v. Group 14 had been a successful group with a large territory (Fig 1). In early September 2002, the group included the pair and six offspring from three different years (Table 1); by mid October, a one-and-a-half-year old male (CC) was the sole survivor. He was observed alone at times in his territory, and did not take part in the battles for ownership of parts of it that began once both of his parents had disappeared. In late November, CC was observed with members of Group 115 in what had been his parents’ territory (Group 115 had apparently won ownership), and he remained with them, there, thereafter. In 2003, if CC contributed to nestling feeding he did so at a very low rate. All of Group 115, including CC, disappeared by mid June; by the end of November, approximately 65% of crows in Stillwater had fallen victim to WNv that year (Caffrey et al. 2005).
vi. In late May 2002, 1-2 weeks after four nestlings fledged from his nest, the unmarked but identifiable male breeder of Group 66 (Table 1) disappeared. By within 1-1.5 weeks later, one of the four fledglings disappeared, and the widowed female’s three-year old son (BD; of a father different than the unmarked individual) returned after having dispersed 1.5 years earlier (he had been seen a few times in the interim, in large flocks outside of Stillwater and as part of a large gathering at the beginning of the 2002 breeding season [Caffrey and Peterson 2015]). BD moved right back into his natal territory and into his mother’s proximity (she was seen allopreening him early after his return), and was initially aggressive toward all three of his juvenile half-siblings (at least two of which begged from him). Son and mother seemed to get closer, literally, as population members began, and continued, to die through September and October. One of the three juveniles (a female) disappeared during October, and in November one of the others (a male) dispersed and immigrated into a group my students and I knew little about (#44, Fig. 1). The third (a male) remained with his mother and half brother as the pair bred together in 2003, and assisted in feeding their nestlings. All three fledglings disappeared by the end of June; the rest of the group were all dead by the end of the summer.
vii. As the weeks of WNv-related death in 2002 came to an end, the survivors of Group 100 (previously Group 333; Table 1) – the female breeder (DZ), her mate (a former group member who had fathered one of three of her fledglings in 2001 [a case of reproductive sharing; KP in Appendix A 1b in Caffrey and Peterson 2015), and two second-year offspring of DZ and her former mate – became very close; the four of them were often seen together, and in close proximity to each other, through the end of 2002 and beginning of 2003. Both two-year olds spent most of their time elsewhere while DZ and KP fed nestlings, but returned at fledgling to incorporate right back into the group. DZ, both two two-year olds, and all three fledglings were dead by mid September. KP survived the Fall of 2003 and by mid October had merged with the few survivors of Group 55; a group having budded from Group 333 (100) in the past (Appendix A 1b in Caffrey and Peterson 2015).

